# Culturomics unveils species and expands bacterial and fungal diversity in Inuit oropharyngeal microbiota

**DOI:** 10.64898/2026.04.20.719640

**Authors:** Mathilde Flahaut, Philippe Leprohon, Nguyen Phuong Pham, Hélène Gingras, Jean Bourbeau, Barbara Papadopoulou, François Maltais, Marc Ouellette

**Affiliations:** Centre de Recherche en Infectiologie et Axe des maladies Infectieuses et immunitaires du Centre de Recherche du CHU de Québec, Canada; Département de Microbiologie, Infectiologie et Immunologie, Faculté de Médecine, Université Laval, Québec City, Québec, Canada; Montreal Chest Institute of the McGill University Health Center; Institut universitaire de cardiologie et de pneumologie de Québec

**Keywords:** Culturomics, oropharyngeal microbiota, mycobiota, *Pauljensenia*, *Shaalia*

## Abstract

Recent advances in high-throughput sequencing and novel culture techniques have revolutionized our understanding of the human microbiota. However, most studies primarily focused on bacterial communities, often overlooking the fungal component. Building upon our previous metagenomic analysis of the Inuit oropharyngeal microbiome ^1^, this study used culturomics to provide a more comprehensive view of both bacterial and fungal communities. We analyzed oropharyngeal swabs from the Qanuilirpitaa? 2017 Inuit Health Survey ^2^, demonstrating the complementarity of metagenomic and culturomic approaches. Our findings highlight the importance of culturomics in revealing low-abundance microorganisms, particularly fungi, which are often underrepresented in metagenomics data. Moreover, we designed an approach to isolate previously uncultivated species. We described two *Pauljensenia* sp., and provided insights into the phylogenetic relationship between *Schaalia* and *Pauljensenia* genera. This study underscores the necessity of a holistic approach to microbiome research, combining multiple techniques to fully elucidate microbial diversity in unique populations like the Inuit.

## Introduction

Several technological advances have greatly enhanced our understanding of microbiota. First, high-throughput sequencing has considerably increased the detection and identification of numerous microorganisms ^3^. The speed and the ability to assess the abundance of a broad range of microbes are the main strengths of this method ^4^. Novel culture techniques, particularly those used for highly stringent species, have revolutionized our concepts of the microbiota ^3,5^. Genome assemblies derived from metagenomic data may lead to incomplete or fragmented genomes and the sequencing of cultured isolates can help in yielding genome sequences of higher quality ^6^. Culturomics consists of a wide variety of culture conditions, and a high-throughput method for colony identification. This enabled the description and classification of several bacteria, archaea, and fungi that had previously been impossible to isolate and study ^6–8^. Recently, the classification of taxa has undergone substantial changes due to the combined efforts of these methods. For example, in 2018, the number of species reported to have been isolated at least once from the human body as pathogens or commensals increased from 2 172 to 2 776 ^9^. Of these novel species, 288 were newly cultured ^9^. The same year, Nouioui *et al*., proposed a new classification of the phylum Actinobacteria with the recognition of 2 orders, 10 families, 17 genera, and the transfer of over 100 species to other genera ^10^. Still, some inconstancies remain among these novel genera, especially between *Pauljensenia* and *Schaalia*, as genomes may be assigned different species names depending on the database used for taxonomic classification. For example, reference genomes annotated as *P. odontolytica* in the Genome Taxonomy Database (GTDB) (gtdb.ecogenomic.org, accessed March 2026) correspond to *S. odontolityca* genomes in the NCBI database. This demonstrates the necessity of culturomic investigations to enhance our understanding of human microbiota. Additional efforts must also be made on mycobiota. Since its launch in 2007, the Human Microbiome Project has made significant advances on the human microbiome but most studies have primarily focused on bacteria, often neglecting Fungi due to their low abundance ^11^.

Metagenomics provides a comprehensive inventory of the DNA of the most abundant species in an ecosystem. Culturomics complements this picture by distinguishing living species and highlighting minority species that may not be correctly or partially detected ^4^. Dealing with low microbial density is especially important in research on the oropharyngeal (OP) microbiota owing to the predominance of host DNA on the OP swabs ^7^. In 2023, we published a metagenomic study on the link between respiratory health in Inuit communities and their airway microbiome ^1^. We showed that Inuit microbiota composition was distinct from other populations, and highlighted differences in taxa or metabolic potential according to the age, sex or respiratory capacities of the participants. As a complement to this previous metagenomics analysis, we report here the analysis of the mycobiota (through metagenomics- and culture-based approaches) and the bacterial culturomics of the same OP swabs obtained as part of the Qanuilirpitaa? 2017 Inuit Health Survey ^2^. We highlight the complementarity of metagenomics and culturomic approaches, especially for the mycobiota. We also describe two previously uncultivated species and provide clarification to the phylogeny of the genera *Schaalia* and *Pauljensenia*.

## Results

### Culture enrichment complements metagenomics in taxonomic identification

This study analyzed the OP microbiota recovered from 180 Inuit that participated in the Qanuilirpitaa? 2017 Inuit Health Survey ^2^. Eighty-five of these participants showed signs of bronchial obstruction, as their forced expiratory volume in one second (FEV1) over forced vital capacity (FVC) ratio was below 0.7, as measured by spirometry. These were matched for age and sex with 95 participants who had normal spirometry (defined as a FEV1/FVC > 0.7 and a FVC > 80%). To conduct an in-depth survey of the OP microbiota composition, OP swabs were processed in two parts; one for metagenomic sequencing ^1^ and the other for culturomics (this study). For the latter, OP swabs were used to seed 16 culture media, 13 for bacteria and 3 for fungi. For bacteria, each medium was inoculated in duplicate, one for incubation under anaerobic conditions and another for incubation under a 5% CO_2_ atmosphere (Figure 1).

**Figure 1.**
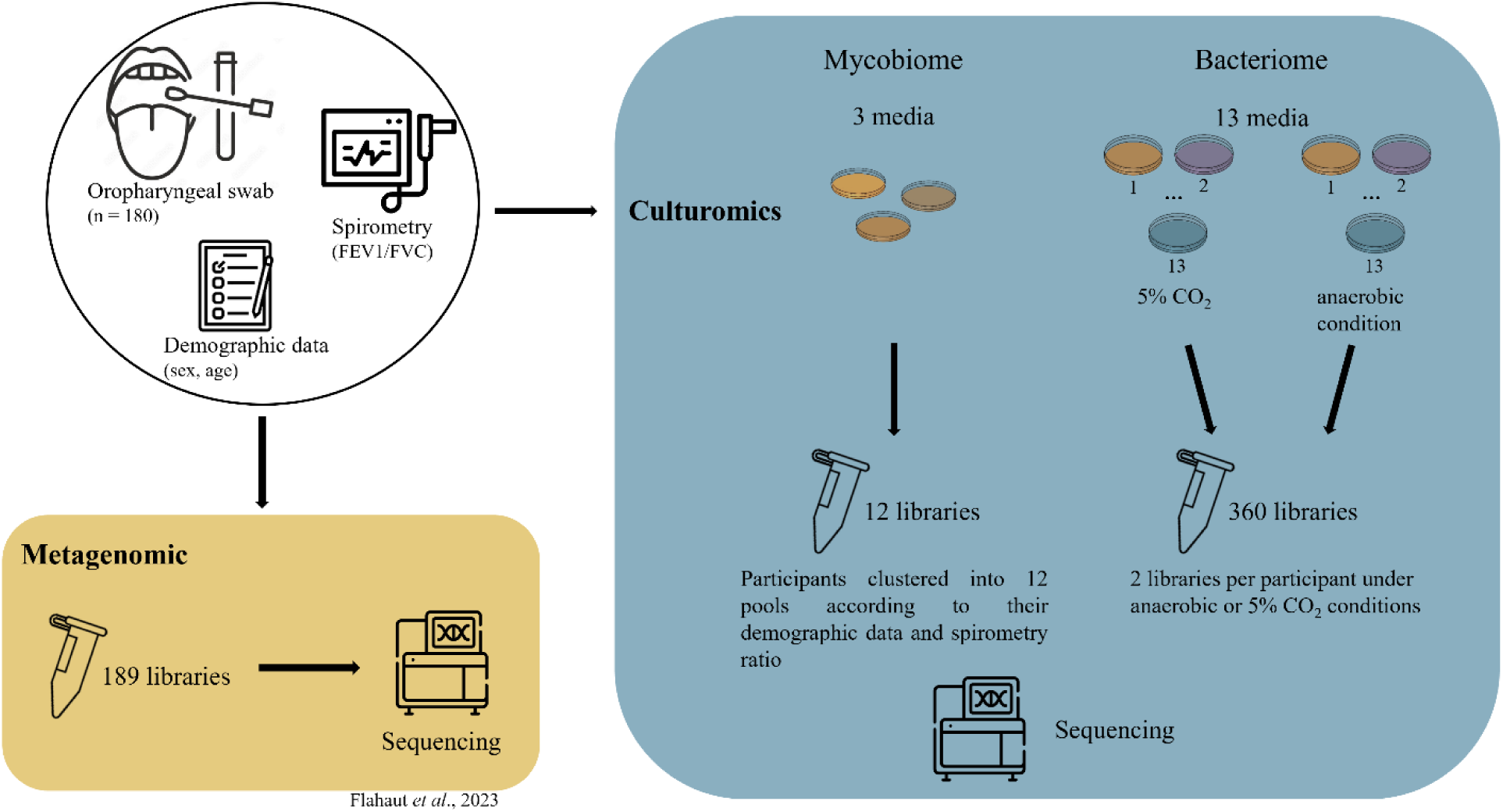
Workflow for the processing of oropharyngeal swabs from Inuit participants. Oropharyngeal swabs from 180 participants were characterized by metagenomic sequencing ^1^ and used to seed 3 culture media for fungi and 13 culture media for bacteria. The latter were seeded in duplicates for incubation in either 5% CO_2_ or anaerobic conditions. For fungi media, participants were categorized into 12 distinct groups according to their demographic data and spirometry ratio (see Supplementary Table 1). For each group, fungal colonies were scrapped from the plates and pooled for sequencing. For bacteria, colonies from the 13 media were scrapped from the plates and pooled in an atmosphere- and participant-wise fashion; for each participant we thus had two pools, one for colonies grown under anaerobic conditions and another for colonies grown under 5% CO_2_.

For the mycobiota analysis, samples were categorized into 12 distinct groups according to the demographic data and spirometry ratio of the participants (Supplementary Table 1). For each group, fungal colonies were scrapped from the plates and pooled for sequencing. Taxonomic profiling of this cultivated mycobiota was performed using Kraken2 (whose database is the most extensive for fungi) and allowed the identification of 22 fungal species from 15 genera (Figure 2A). In comparison, only 8 fungal species (from 6 genera) were detected among the groups when taxonomic profiling was performed from metagenomic sequencing of the swabs (Figure 2A). Of these, 5 species (from 4 genera) were common to both methods (Figure 2A), including *C. albicans*, *C. dubliniensis*, and *Nakaseomyces glabratus* (previously part of the *Candida* genus), which were also among the most prevalent between the groups of participants (Figure 2B). Our data indeed indicates that metagenomics identified species highly prevalent among the groups while culturomics enabled also the identification of fungal species with a sparser distribution. Still, some genera were detected solely by metagenomics (*Akanthomyces* and *Aspergillus*) or were poorly detected by fungal culturomics (*Malassezia*) (Figure 2B), which may relate to the culture media used.

**Figure 2.**
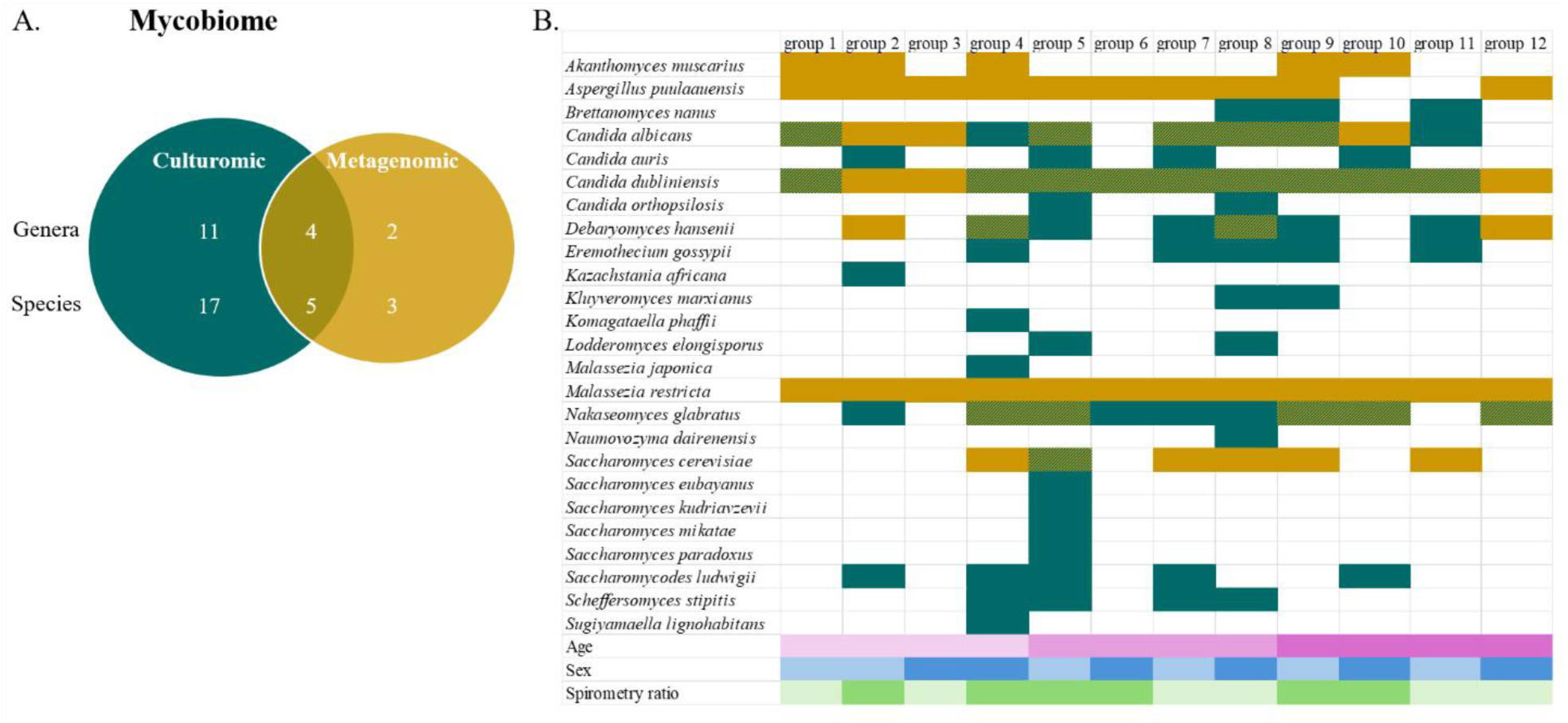
Fungal genera and species identified by Kraken2 from metagenomic and culturomic reads. (A) Venn diagram of the number of genera and species identified for each approach. (B) HeatMap for the presence of fungal species in culturomics (dark green), metagenomics (yellow) or both (green) datasets among the 12 groups of participants (see Supplementary Table 1) defined by the age (light pink, 16-35 y.o.; medium pink, 36-55 y.o.; dark pink, 56-71 y.o.), sex (light blue, Male; dark blue, Female) and spirometry ratio (light green, >0.7; dark green, <0.7) of the participants.

For bacteria, colonies from the 13 media were scrapped from the plates and pooled in an atmosphere- and participant-wise fashion; for each participant we thus had two pools, one for colonies grown under anaerobic conditions and another for colonies grown under 5% CO_2_ (Figure 1). We noted that blood agar media (CBA, CNA, and PEA) as well as TSY, BHI, and CHOC (for abbreviations see Materials and Methods section and Supplementary Table S2) were the media yielding the highest colony counts regardless of oxygenation conditions. Conversely, MAC and MSA were the media with the lowest growth and sometimes remained completely sterile, which may be due to their selectivity (Supplementary Table 2). Taxonomic profiling of the participants’ pools of colonies using MetaPhlAn4 revealed a total of 125 bacterial genera and 361 species (Figure 3A). In comparison, the previous metagenomic sequencing conducted from the same samples ^1^ yielded 362 bacterial genera and 933 species (for proper comparison with culturomics this previous metagenomic data was re-analyzed here with MetaPhlAn4 as only MetaPhlAn3 was available at the time of the initial publication. Substantial overlap was noted between the two approaches and most species (and genera) detected by metagenomics included those found by culturomics (Figure 3A). The major phyla were also similar between culturomics (Figure 3B) and metagenomics ^1^, with the Firmicutes, Actinobacteria, Proteobacteria, and Bacteroidota dominating. However, minor variations were observed in the ranking of the most prevalent genera when comparing the two methods. Indeed, while culturomics allowed a better representation for the genera *Aggregatibacter*, *Lacticaseibacillus*, and *Staphylococcus*, it underperformed compared to metagenomics for the genera *Actinomyces*, *Granulicatella*, and *Lancefieldella* (Supplementary Figure 1). For *Staphylococcus* the difference further extends to the number of species identified, with only 8 *Staphylococcus* species detected by metagenomics while 16 were found by culturomics. In contrast, *Rothia*, *Schaalia*, *Streptococcus,* and *Veillonella,* which were among the most abundant genera in our samples, were detected at very similar occurrences by both approaches (Supplementary Figure 1). Of the 10 genera and 34 species detected specifically by culturomics, *Heyndrickxia* (and specifically the species *H. sporothermodurans*) was the most prevalent and was detected in 10% of the samples (Figure 3C). *Brevibacillus* was the second most prevalent culturomic-specific genus and was detected in 11 samples, with its species *B. agri* detected in 8 samples (Figure 3C).

**Figure 3.**
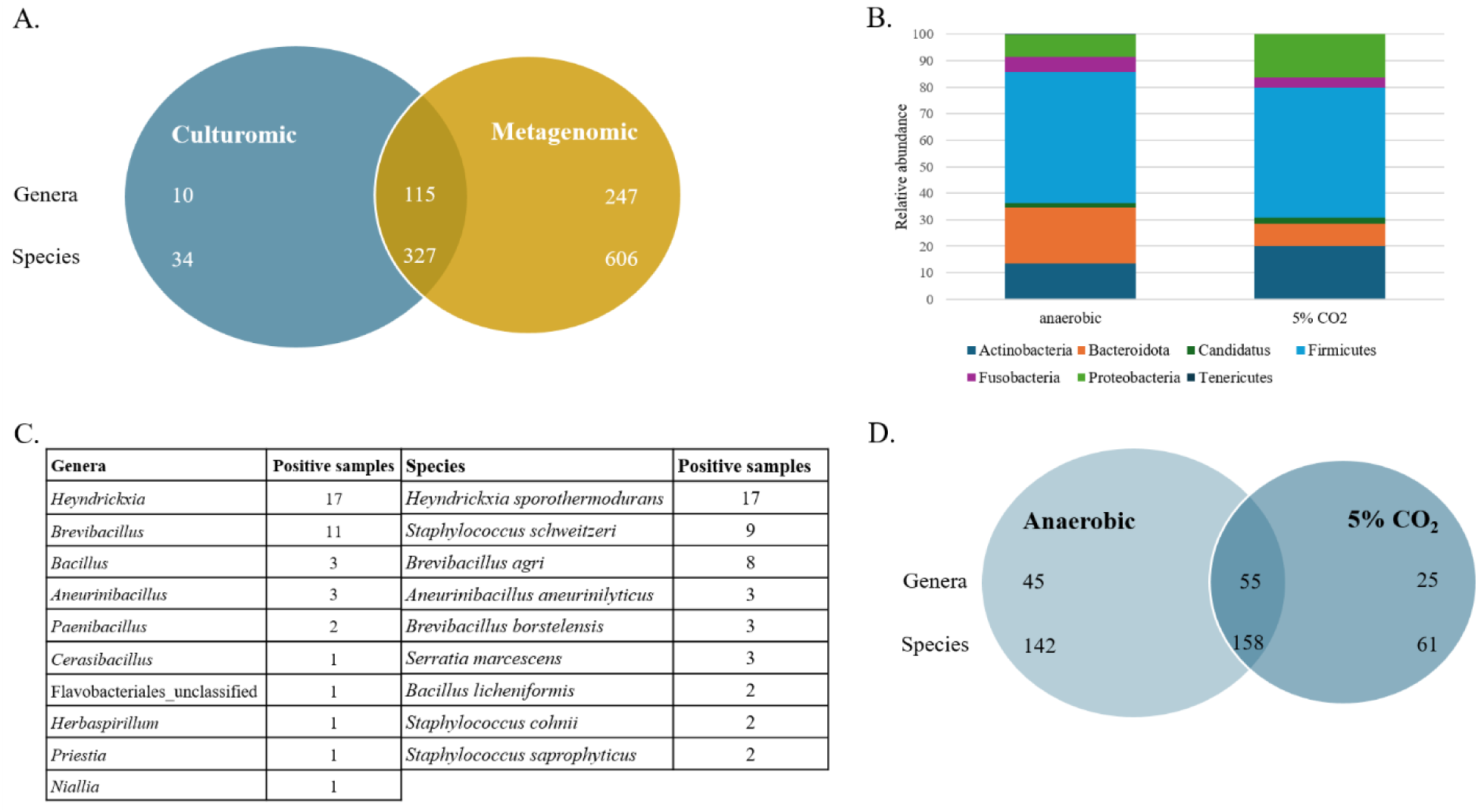
Bacterial genera and species identified by MetaPhlAn4 from metagenomics and culturomic reads. (A) Venn diagram of the number of genera and species identified for each approach. (B) Relative abundance of culturomics identified phyla under 5% CO_2_ and anaerobic conditions. (C) The bacterial genera and species identified specifically by culturomics. For the species, only those found in more than 2 samples are shown. (D) Venn diagram of the number of genera and species recovered from plates incubated under 5% CO_2_ or anaerobic conditions.

Lastly, the use of aerobic and anaerobic conditions allowed the identification of substantially more species/genera than either condition alone (Figure 3D). Overall, culturomics identified 125 bacterial genera and 361 species (Supplementary Table 3, sheet 1). Among these, 25 genera and 61 species were found specifically in cultures incubated in 5% CO_2_ whereas 45 genera and 142 species were classified as strictly anaerobic (Figure 3D). Incubation under aerobic conditions yielded more Proteobacteria and Actinobacteria, at the expense of Bacteroidota and Firmicutes (Figure 3B). Indeed, 12 of the 27 genera strictly identified in 5% CO_2_ belonged to Proteobacteria; of the 44 genera exclusively identified in anaerobic cultures, 19 and 12 were part of Firmicutes and Bacteroidota, respectively (Figure 3D and Supplementary Table 3, sheet 2 and 3). At the species level, anaerobic cultures yielded exclusively 22 of the 29 *Prevotella* species identified in our samples (Supplementary Table 3, sheet 4). Conversely, of the 17 *Staphylococcus* species retrieved, 12 were exclusive to cultures incubated under 5% CO_2_ whereas only 3 were strictly identified in anaerobic conditions (Supplementary Table 3, sheet 5).

Overall, our data highlights the complementary nature of 5% CO_2_ and anaerobic culturing techniques in capturing the full diversity of cultivable bacterial species. Moreover, our comparison of metagenomics and cultured methods suggests that while the approaches converge on dominant taxa, they offer complementary insights into the less abundant microbial community members.

### Differences in cultured profiles occur based on demographic data and respiratory capacities of the participants

We next examined whether the occurrence of fungal or bacterial species (or genera) differed among participants based on demographic characteristics (age or sex) or spirometry ratios. For fungal cultures, the 36-55-year-old (y.o.) participants demonstrated the highest level of diversity with 18 distinct fungal species identified, whereas 12 and 9 species were recovered from the 16-35 and 55-71 y.o. participants, respectively (Figure 2B). Of note, the species *Candida orthopsilosis*, *Lodderomyces elongisporus*, and *Scheffersomyces stipites* were almost exclusively observed in pools grouping participants aged 36-55 y.o., while these were specifically devoid of *Akanthomyces muscarius* (Figure 2B). The summed counts of culturomic-derived species were also twice as much for groups of participants aged 36-55 y.o. (n=35) compared to groups derived from participants aged 16-35 y.o. (n=16) or 56-71 y.o. (n=17) (Figure 2B), a large part being attributed to 36-55 y.o. male suffering from bronchial obstruction (group 5). Globally, participants with bronchial obstruction (FEV1/FVC < 0.7) showed a higher occurrence of *Nakaseomyces glabratus*, *Saccharomycodes ludwigii*, and *Candida auris* compared to control participants (FEV1/FVC > 0.7) (Figure 2B). *Debaryomyces hansenii, Candida albicans,* and *Candida auris* were slightly more common in culturomic pools of male participants than in pools of females (Figure 2B).

For bacterial cultures, differences in the distribution of bacterial species (or genera) were found according to the age, sex and respiratory function of participants (Figure 4). It is important to note that for each comparison described below, bacterial genera or species found at a prevalence <5% among samples of a given category were excluded from this analysis. When looking for differences based on age, we found that the genera *Olsenella* and *Shuttleworthia*, with their species *Olsenella phocaeensis* and *Shuttleworthia satelles*, were more prevalent among the youngest participants (16-35 y.o.) (Figure 4A). The same was true for the species *Corynebacterium durum*, *Prevotella oris*, and *Veillonella* sp CHU110 (Figure 4A). For *Prevotella oris*, the increased prevalence among participants aged 16-35 y.o. had also been found by metagenomics ^1^. The species *Alloprevotella* SGB1463, *Capnocytophaga sputigena* (the only species more prevalent among participants with bronchial obstruction (Figure 4B)), and *Haemophilus parainfluenzae* were also more prevalent among participants aged 16-35 y.o., but only compared to 36-55 y.o. participants, while the species *Parvimonas* sp oral taxon 110 and *Veillonella* sp 3627 were specifically more prevalent compared to participants aged 56-71 y.o. (Figure 4A). The species *Atopobium* sp oral taxon 199 and *Oribacterium* SGB96486 were absent from the 16-35 y.o. participants (Supplementary Table 3, sheet 6) while the oldest participants (56-71 y.o.) specifically lacked the species *Dolosigranulum pigrum, Eubacterium infirmum*, and *Mogibacterium* SGB3923 (Supplementary Table 3, sheet 6), but had a higher prevalence of *Neisseria bacilliformis* compared to 16-35 y.o. participants (Figure 4A).

**Figure 4.**
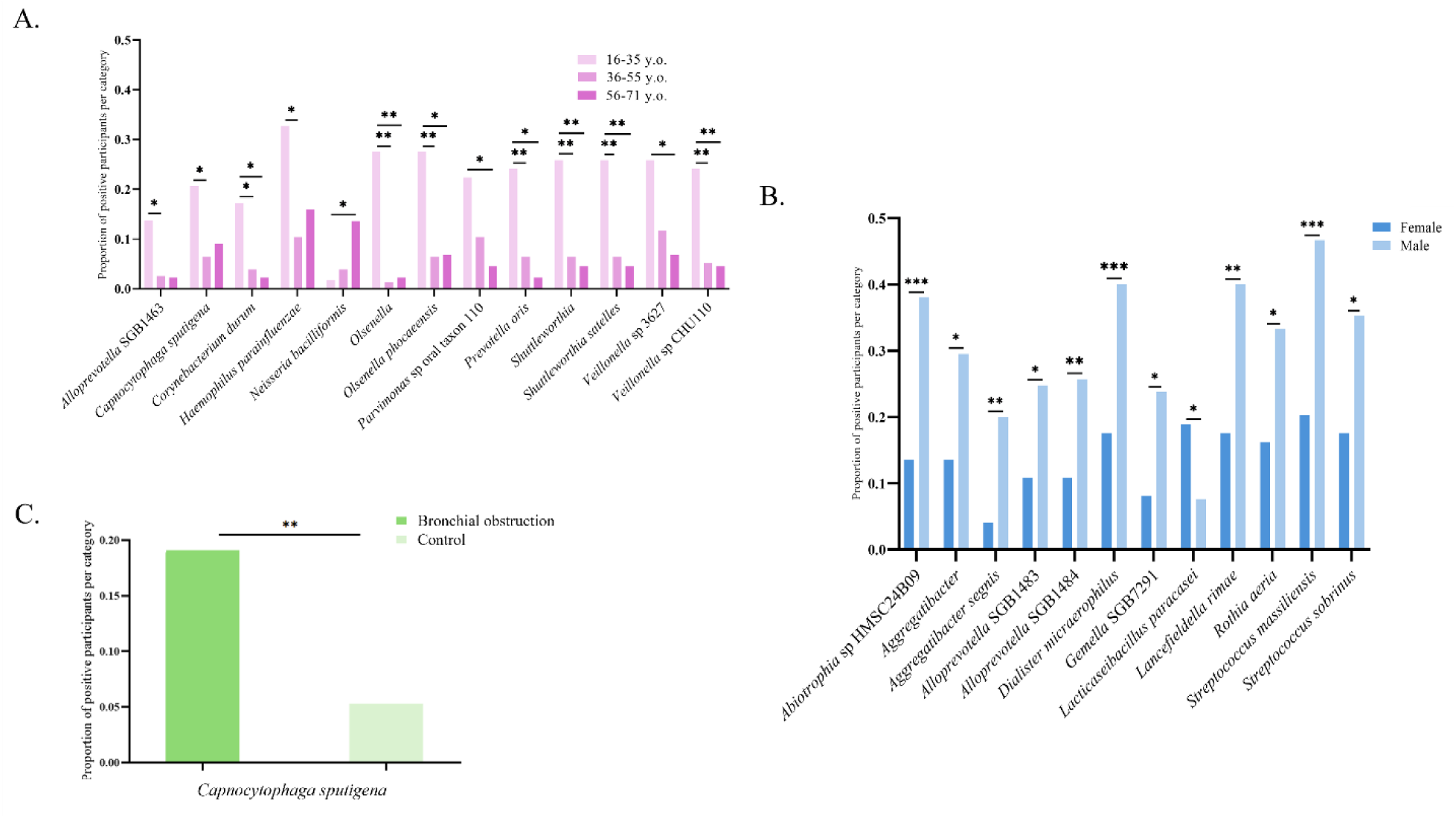
Cultured bacterial taxa identified by MetaPhlAn4 and associated with the demographic data or spirometry ratio of the participants. Bacterial genera and species whose prevalence differs according to the age (A), sex (B) or respiratory health status (C) of the participants. Chi-square test was performed on genera and species at least twice as prevalent in one group, per category.

Regarding differences according to sex, the genus A*ggregatibacter*, with its species *A. segnis*, was twice as much prevalent among male participants than female participants (Figure 4C). The same is true for eight other species, including *Streptococcus massiliensis* and the unnamed species *Abiotrophia* sp HMSC24B09, *Alloprevotella* SGB1483, *Alloprevotella* SGB1484, and *Gemella* SGB7291, which all had an increased prevalence among male participants (Figure 4C). Interestingly, *S. massiliensis* was also shown to be more prevalent among male by our previous metagenomic study ^1^. The genus GGB38873 (with its species GGB38873 SGB47522), and the species *Oribacterium* SGB96486 and *Neisseria cinerea* were identified only in male participants (Supplementary Table 3, sheet 6). The only species more prevalent among female participants was *Lacticaseibacillus paracasei* (Figure 4C).

### Culturomics allows isolation and description of previously uncultured species

For our investigation of poorly characterized microbial taxa, we focused our analysis on genera with unnamed species. No novel or uncultivated fungal species were revealed with Kraken2 so we focused our work on bacteria. We first reconstructed genomes from the bacterial culturomic data and processed these with GTDB-Toolkit (GTDBTk) for taxonomic profiling. We succeeded in reconstructing 852 genomes with a completeness > 90% and contamination < 5% (Supplementary Table 4, sheet 1). These included 171 different species across 52 genera (Figure 5).

**Figure 5.**
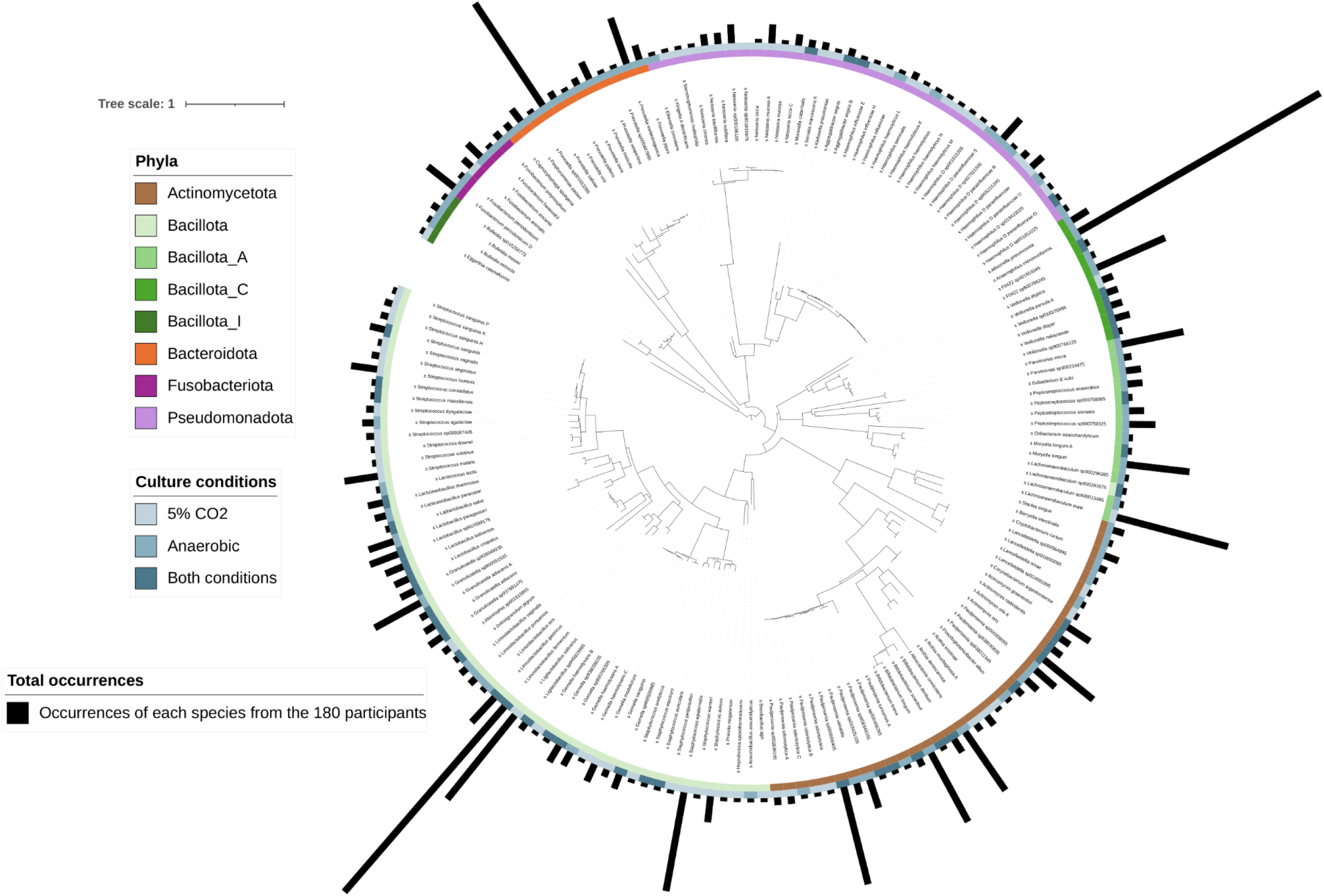
Phylogenetic tree constructed from the genome of the 171 bacterial species assembled from culturomic data. Genomes were classified by GTDB-Tk v2.4.0 and the phylogenetic tree was constructed with PhyloT (phylot.biobyte.de) and visualized in iTOL. The phyla of the genomes are shown by the inner circle and the color scheme indicated in inset. Genomes assembled from bacteria grown specifically under 5% CO_2_ (light blue) or anaerobic condition (medium blue), and those grown under both atmosphere (dark blue) are shown by the outer circle. The length of the outer black bars indicates the relative prevalence of the species among the samples, minimum n = 1, maximum n = 75.

The genera *Limosilactobacillus, Pauljensenia* and *Anaeroglobus* were detected most frequently and were found in 80, 79, and 75 samples, respectively (Supplementary Figure 2 and Supplementary Table 4, sheet 2). *Pauljensenia* sp900556405 was the most frequently observed species with 27 positive samples from both 5% CO_2_ and anaerobic conditions (Table 1). Forty-seven cultures had genomes that could not be assigned at the species level, *Pauljensenia* dominated with 18 unnamed species, followed by *Gemella* with 12 unnamed species, and by *Granulicatella* and *Alloscardovia* each with 5 unnamed species (Table 1 and Supplementary Figure 3).

**Table 1.** Counts of samples (by atmospheric conditions) and participants positive for bacterial genera without a species name.

| **Taxons<sup>a</sup>** | **5% CO<sub>2</sub>** | **Anaerobic** | **Total number of positive cultures** | **Number of participants with positive culture(s)<sup>b</sup>** |
| --- | --- | --- | --- | --- |
| ***Pauljensenia*** | 29 | 41 | 70 | 49 |
| *Pauljensenia* sp | 8 | 10 | 18 | 15 |
| *Pauljensenia* sp000308055 | 3 | 4 | 7 | 7 |
| *Pauljensenia* sp000466265 | 1 | 0 | 1 | 1 |
| *Pauljensenia* sp001838165 | 0 | 2 | 2 | 2 |
| *Pauljensenia* sp019425105 | 4 | 4 | 8 | 7 |
| *Pauljensenia* sp900556405 | 11 | 16 | 27 | 23 |
| *Pauljensenia* sp938022345 | 0 | 2 | 2 | 2 |
| *Pauljensenia* sp938030835 | 2 | 2 | 4 | 3 |
| *Pauljensenia* sp958349165 | 0 | 1 | 1 | 1 |
| ***Granulicatella*** | 12 | 15 | 27 | 23 |
| *Granulicatella* sp | 1 | 4 | 5 | 5 |
| *Granulicatella* sp900551535 | 6 | 8 | 14 | 11 |
| *Granulicatella* sp916049935 | 2 | 2 | 4 | 3 |
| *Granulicatella* sp937891475 | 3 | 1 | 4 | 3 |
| ***Gemella*** | 17 | 5 | 22 | 21 |
| *Gemella* sp | 10 | 2 | 12 | 11 |
| *Gemella* sp900555985 | 2 | 2 | 4 | 4 |
| *Gemella* sp900766305 | 4 | 1 | 5 | 5 |
| *Gemella* sp938038035 | 1 | 0 | 1 | 1 |
| ***Haemophilus*** | 7 | 6 | 13 | 12 |
| *Haemophilus* sp | 1 | 0 | 1 | 1 |
| *Haemophilus*_D sp001811025 | 2 | 0 | 2 | 2 |
| *Haemophilus*_D sp019423025 | 1 | 0 | 1 | 1 |
| *Haemophilus*_D sp905215245 | 0 | 2 | 2 | 2 |
| *Haemophilus*_D sp927911595 | 3 | 4 | 7 | 6 |
| ***Peptostreptococcus*** | 0 | 8 | 8 | 8 |
| *Peptostreptococcus* sp000758885 | 0 | 1 | 1 | 1 |
| *Peptostreptococcus* sp900759325 | 0 | 7 | 7 | 7 |
| *Lactobacillus* sp910589175 | 2 | 6 | 8 | 8 |
| **F0422** | 0 | 6 | 6 | 6 |
| F0422 sp | 0 | 4 | 4 | 4 |
| F0422 sp001553345 | 0 | 1 | 1 | 1 |
| F0422 sp900766245 | 0 | 1 | 1 | 1 |
| *Alloscardovia* sp | 0 | 5 | 5 | 5 |
| *Neisseria* sp000186165 | 5 | 0 | 5 | 5 |
| ***Lancefieldella*** | 0 | 4 | 4 | 4 |
| *Lancefieldella* sp000564995 | 0 | 2 | 2 | 2 |
| *Lancefieldella* sp024691995 | 0 | 1 | 1 | 1 |
| *Lancefieldella* sp916050095 | 0 | 1 | 1 | 1 |
| ***Veillonella*** | 3 | 1 | 4 | 4 |
| *Veillonella* sp018376995 | 2 | 1 | 3 | 3 |
| *Veillonella* sp900766125 | 1 | 0 | 1 | 1 |
| ***Streptococcus*** | 3 | 0 | 3 | 3 |
| *Streptococcus* sp | 1 | 0 | 1 | 1 |
| *Streptococcus* sp000187445 | 2 | 0 | 2 | 2 |
| ***Lachnoanaerobaculum*** | 0 | 3 | 3 | 3 |
| *Lachnoanaerobaculum* sp000287675 | 0 | 1 | 1 | 1 |
| *Lachnoanaerobaculum* sp000296385 | 0 | 1 | 1 | 1 |
| *Lachnoanaerobaculum* sp938015485 | 0 | 1 | 1 | 1 |
| *Parvimonas* sp000214475 | 0 | 2 | 2 | 2 |
| ***Prevotella*** | 0 | 2 | 2 | 2 |
| *Prevotella* sp000467895 | 0 | 1 | 1 | 1 |
| *Prevotella* sp001553265 | 0 | 1 | 1 | 1 |
| *Abiotrophia* sp001815865 | 1 | 1 | 2 | 2 |

<sup>a</sup> Taxonomic profiling made with GTDB-Tk v2.4.0.
<sup>b</sup> The number of participants with positive culture(s) can be different from the total number of
positive samples when the species is found both in 5% CO<sub>2</sub> and anaerobic conditions.

The diversity observed for the genus *Pauljensenia*, with a large number of unnamed species (Supplementary Table 4, sheet 3), was particularly noteworthy as it had not been detected while profiling the culturomic sequencing reads using MetaPhlAn4, which instead led to a high number of unnamed species from the genus *Schaalia* (Supplementary Table 3, sheet 7). This led us to note that the classification of species within the *Pauljensenia* genus varies depending on the database used. *Pauljensenia* species in GTDB include mostly species referred as *Schaalia* and *Actinomyces* in other databases, while the GTDB does not contain any genomes referred to as *Schaalia* (gtdb.ecogenomic.org, accessed March 2026). This discrepancy illustrates the variability in taxonomic nomenclature across different databases. We wanted to address the limited knowledge surrounding this relatively understudied *Pauljensenia* genus, which was first described in 2018 ^10^, by isolating *Pauljensenia* species from our culturomic material. We thus compared through reciprocal BLAST the *Pauljensenia* genomes reconstructed from our culturomic data, the *Pauljensenia* genomes from GTDB, the *Schaalia* genomes from NCBI, as well as a number of outlier genomes to identify a gene that could be amplified by PCR specifically from *Pauljensenia*. The BLAST output was used to compute the mean gene percentage identity of the samples compared to the *Pauljensenia* genome GCA_001064145.1 which was used as reference for the reciprocal BLAST. From these values, we could easily observe two different populations whether the genomes came from our culturomic data, from GTDB or from NCBI database (Supplementary Figure 4). We identified the gene JVOJ01000036.1_32 (Supplementary figure 6) as uniquely found in the genomes from the group harboring the greatest proportion of unnamed *Pauljensenia* species within our samples and designed PCR primers for screening small round colonies derived from three samples (729a, 757a and 895a). We obtained ten positive colonies from the three participants and sequenced their genomes. Colonies were translucent to white with regular edges on blood agar plates after incubation for 48 h at 35 °C under 5% CO_2_ condition.

The ten reconstructed genomes demonstrated high quality, as evidenced by their low contamination levels, great completeness (with a minimum of 92.22%), and few mismatches or ambiguous bases (Supplementary Table 5, sheet 1). The GC content ranged from 64.9% to 66.9% (Supplementary Table 5, sheet 1) and all isolates were capnophilic facultative anaerobes, Gram positive with a cell size of 0.5–0.9 x 0.5–1.3 µm (Figure 7). Two were classified by GTDB-Tk as *Pauljensenia odontolytica*, three as *Pauljensenia* sp958349165, and five as *Pauljensenia* sp900556405 (Figure 6 and Supplementary Table 5, sheet 1). Our genome reconstructions yielded superior results compared to the uncultured *Pauljensenia* reference reconstructions available on GTDB (Supplementary Table 5, sheet 2). The examination of the three species using an electron microscope revealed polymorphism with a high propensity for division, as they were observed in the logarithmic phase. Notably, these bacteria exhibited Y-shaped division patterns, budding, binary division, and atypical cell division specific to the Actinomycetaceae family (Figure 7) ^12^. We did not notice any flagellum or spore.

**Figure 6.**
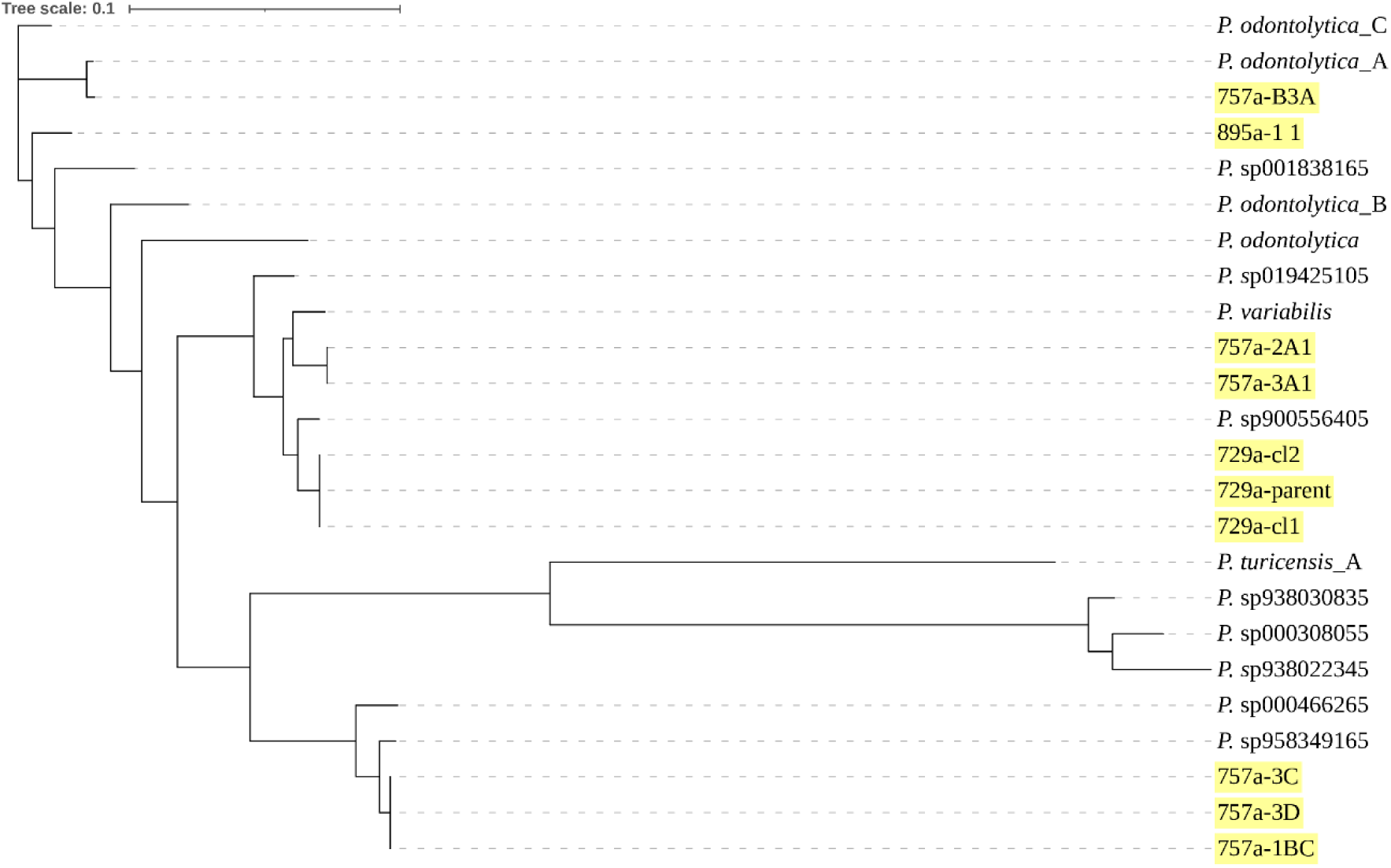
Phylogenetic tree of reconstructed *Pauljensenia* genomes. Genomes were annotated using PROKKA v1.14.5, clustering with Panaroo v1.0.1, aligned with mafft v7.526, the phylogenetic tree was constructed with FastTree v2.1, and visualized using Dendroscope 3 v3.8.10. The tree was built from the 10 *Pauljensenia* colonies isolated on plates (highlighted in yellow) and the fourteen GTDB species identified by GTDB-Tk v2.4.0 from our culturomic sequencing.

**Figure 7.**
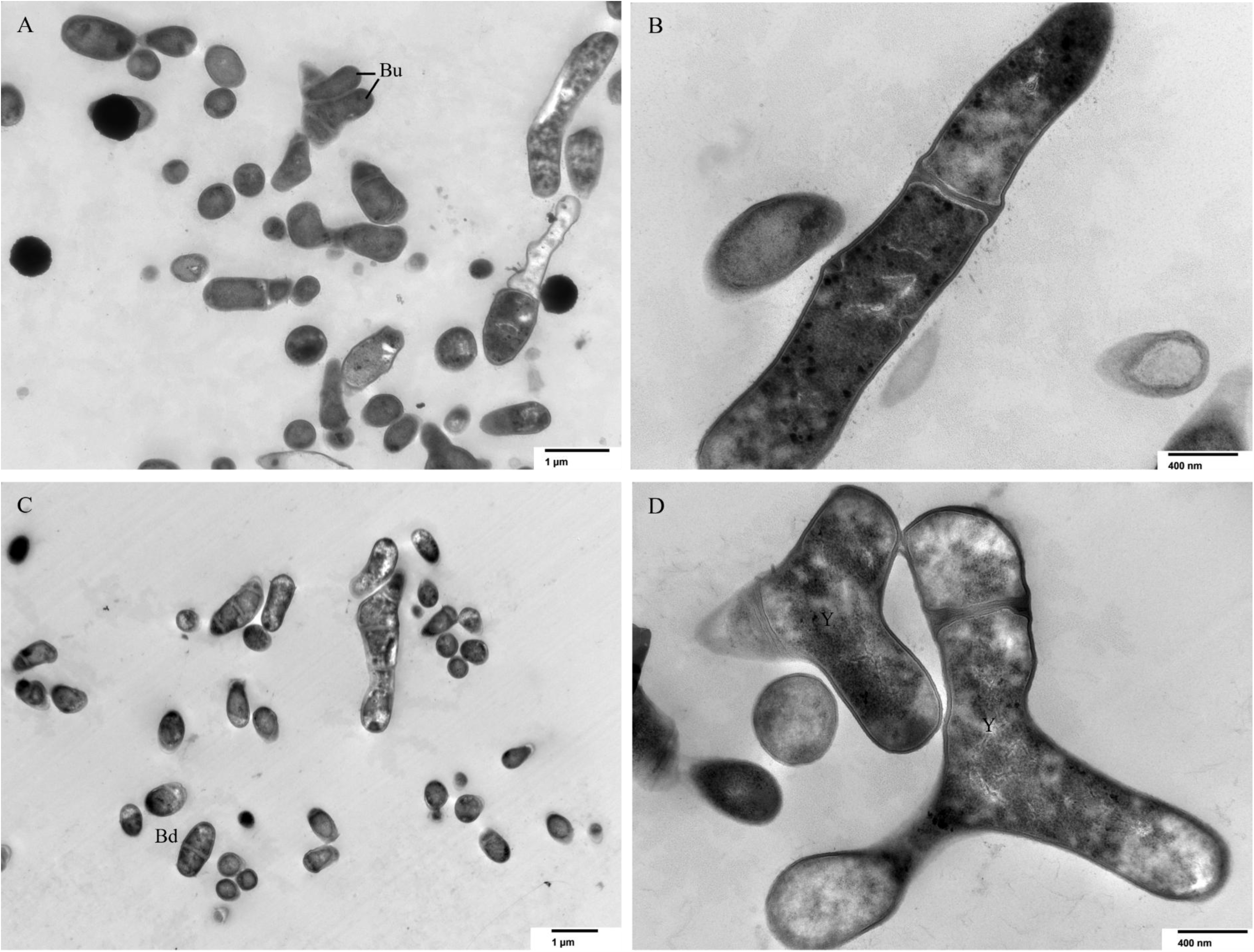
Morphological diversity of *Pauljensenia* species revealed by transmission electron microscopy. Pictures showed budding (Bu), binary division (Bd), Y-shaped division patterns (Y), and atypical cell division. Exposure: 4000 (ms); gamma: 1.00, no sharpening, accel. Voltage = 80 kV. (A) and (B) isolate 757a-1BC (*P.* sp958349165); (C) and (D) isolate 757a-3A1 (*P.* sp900556405).

## Discussion

Metagenomics showed that a large fraction of bacteria remains uncultured. Culturomics, as a complementary approach, allows to maximize the characterization of microbial species from a given ecosystem by increasing the number of cultivated bacteria and fungi ^7^. In the context of the respiratory tract, culturomics has experienced significant growth in recent years, but have often focused on a limited number of culture media or samples ^3,6,13^. Our study, which complement our previous metagenomic characterization of Inuit oropharyngeal samples ^1^, thus expands the repertoire of cultured throat microbiota by the systematic sequencing of commensal bacteria and fungi across a total of 180 samples cultured on 16 different growth media.

In the respiratory tract, few studies have focused on the mycobiota, notably because it represents a minor component of the microbiota ^8^. Our data show that the Inuit mycobiota is dominated by *Aspergillus*, *Candida*, and *Malassezia*, which were the most frequently observed genera both by metagenomics or culturomics. This is consistent with the most abundant genera described in other metagenomic studies for the respiratory tract and oral mycobiota ^14–18^, and with *Candida* species having been shown to colonize 30% to 60% of healthy adults ^19–21^. For the latter genus, our culturomic approach was useful in recovering species with a sparser distribution among participants. Similarly, *Saccharomyces*, a genus frequently detected in the respiratory tract or oral cavity ^14,15,17^, was detected only in a few samples. It was shown that the airway mycobiota in individuals of Asian ascent was dominated either by *Candida* or *Saccharomyces* but rarely by both ^15^, and it is possible that the same happens here for Inuit with *Candida*-dominant profiles. Of note, the detection of *Nakaseomyces* (previously named *C. glabrata*) in most groups of participants may be of concern due to its association with increased resistance to antifungals and hospitalization ^20,22,23^. Several fungal genera detected in our samples were represented by only one species. This aligns with previous findings that revealed a diversity of genera, but with many represented by a single specie ^23^. Such pattern highlights the need for a deeper understanding of fungal diversity, in the oropharynx but also in other niches, to capture the full spectrum of the mycobiota. The isolation of numerous fungal taxa unnoticed by metagenomics demonstrates the power of culturomics in exploring the fungal community structure, and incorporating a broader array of culture media, or additional growth conditions, may uncover an even larger set of rare or novel fungal species with specific growth requirements.

Our culturomic data revealed differences in the bacterial composition of the oropharyngeal microbiota based on age, sex and respiratory health status of the participants. *Capnocytophaga sputigena* was the only species with an increased prevalence among participants with impaired lung function. This species has been implicated in rare cases of pulmonary infections ^24^, mostly in patients with comorbidities, and was also shown to be part of the 4 bacteria whose relative abundance consistently increase across the different stages of gastric carcinogenesis ^25^. A number of species were found to be more prevalent in younger participants (16-35 y.o.), including *Prevotella oris*, *Shuttleworthia satelles*, *Veillonella* sp CHU110, and *Corynebacterium durum*. In contrast, only *Neisseria bacilliformis* was more prevalent in older participants (56-71 y.o.). Similarly, 9 species were more frequent in male participants, while only one was more prevalent in females. This is consistent with the increased bacterial richness observed in younger participants and in male participants by our previous metagenomic sequencing of the same oropharyngeal samples ^1^ and with other studies that showed sex to be a dominant variable in the clustering of respiratory tract metagenomics samples ^26,27^. Such concordance with metagenomics strengthens the validity of our culturomic results. Our study also revealed bacteria exclusively found through culture-based methods, such as *Heyndrickxia* and *Brevibacillus* which were unique to culturomics despite being present in 10% and 6% of samples, respectively. The former includes a species known for its probiotic properties that is currently being investigated for its potential benefits against respiratory symptoms ^28^. Such an example, and that culturomics allowed a better species representation for some genera (e.g. *Staphylococcus*), underscores the importance of tailored culture approaches as a complement to metagenomics in microbiome studies. The choices of growth media and conditions are likely to influence the scope of such approaches, as shown by recent culturomic efforts, including the pioneering Cultivated Oral Bacteria Genome Reference catalog ^29^ and a survey of the airway commensal bacteria ^30^, which were dominated by *Streptococcus* species. In the current study, the 26 growth conditions used for bacteria (13 media incubated under anaerobic and 5% CO_2_ atmosphere) enabled us to reconstruct the genome of 171 species across 52 genera whose relative abundance were rather balanced with no exceedingly abundant genus (Supplementary Figure 5). Also, the oropharynx is a complex ecosystem that harbors a diverse array of microorganisms including anaerobic bacteria ^6,8,29^ and the use of anaerobic growth conditions indeed allowed to broaden bacterial diversity.

Among all genera, *Pauljensenia* was notable for the number of genomes assembled and in having the highest number of genomes of unnamed species. This genus is also an example of the complexity that can arise when profiling microbiome datasets with different genomic resources, most genomes annotated as *Pauljensenia* with GTDB-Tk being annotated as *Schaalia* by the NCBI. GTDB relies on a genome-centric approach, utilizing average nucleotide identity (ANI) and concatenated protein phylogenies for species delineation, whereas NCBI taxonomy relies more heavily on traditional phenotypic and 16S rRNA gene-based classifications. These divergent strategies can lead to inconsistent taxonomic assignments ^31^, as in the case of Actinomycetes (to which *Pauljensensia*/*Schaalia* belongs). Indeed, the *Actinomyces* genus underwent a significant reclassification in 2018, resulting in the creation of several new genera and species, including *Schaalia* sp., and *Pauljensenia* sp. ^10^. With this reclassification, the *Pauljensenia* genus currently comprises only one validated species, namely *P. hongkongensis*, in the List of Prokaryotic Names with Standing in Nomenclature and the *Schaalia* genus encompasses 13 species, with *S. odontolytica* designated as the type species. Following its recent taxonomic reclassification, the *Pauljensenia* genus has emerged as a significant component of the oral microbiome, demonstrating key roles in modulating respiratory health and interactions with human physiological systems ^29,32–34^. In our study, the reconstructed *Pauljensenia* spp. genomes were more likely to come from control participants than from participants with bronchial obstructions. A number of *Pauljensenia* genomes from cultured oral cavity isolates have been recently published ^29^ but for the most part these are distinct from the 3 species isolated and sequenced here. Our genomes are also of higher quality than those of the corresponding GTDB genomes and will thus be useful to refine this newly described genus. This study expands our knowledge, by electronic microscope observations, of this genus which was previously known only for its rod-shape ^10^.

In conclusion, we have applied culturomics to an extensive array of oropharyngeal samples which helped advance our understanding of this complex microbiota. The results presented here highlight the complementary nature of two methodological approaches, metagenomics and culturomics, in microbial community analysis ^7^. Moreover, this comprehensive approach, with the inclusion of the fungal component, has yielded novel insights into the microbial diversity of the oropharyngeal mucosa. Notably, our study led to the isolation and characterization of members of the genus *Pauljensenia*, which, despite its frequent detection in oral and pulmonary mucosal environments, comprises very few cultivated species. This work not only expands our knowledge of oropharyngeal microbiomes but also demonstrates the efficacy of culturomics in elucidating the composition of challenging microbial communities.

## Materials and Methods

### Sampling and culture enrichment

Ethics statement and sampling were carried out in the same way as previously described ^1^. Throat swabs from 180 participants were homogenized and subdivided for distinct analytical protocols. For culture enrichment, 400 µl were diluted 1:10 then 10 µl of the dilution was inoculated per medium. Drawing from the studies carried out by the Surette laboratory ^3^, 13 media for bacteria and 3 media for fungus were used. More details about these selective, non-selective and enriching media are given in Supplementary Table 2. For bacteria, media included McKay medium (MK), mannitol salt agar (MSA, BD), MacConkey agar (MAC, BD), fastidious anaerobe broth with 1,5% agar (FAA, Neogen), actinomycetes isolation agar with 5 ml/L glycerol (AIA, Sigma-Aldrich), brain heart infusion agar (BHI, BD), cooked meat medium (Beef, OXOID), tryptone soya yeast extract agar (TSY, Sigma-Aldrich), Columbia blood agar base (CBA, BD), Columbia CNA agar (CNA, BD), phenylethyl alcohol agar (PEA, BD), GC powder (BD) with 2% haemoglobin and 1% enrichment IsoVitaleX (CHOC, BD), and tryptone soya yeast extract agar with 0.05 µg/L haemin, 10 mg/L vitamin K1, 0.1 mg/L kanamycin, 7.5 mg/L vancomycin, and 5% laked horse blood (KVLB). To CBA, CAN, and PEA 5% of sheep’s blood was added. To Beef, BHI, and TSY, the following additional additives were included: 10 µg/L haemin, 1.0 µl/L vitamin K1, 10 mg/L colistin sulfate, and 0.5 g/L l-cysteine (Supplementary Table 2). Media were inoculated at 35 °C, 24h in 5% CO_2_ conditions and 48h in anaerobic conditions (85% N, 10% H, 5% CO_2_). For fungi, the 3 media were sabouraud 4% glucose agar with 0.4 mg/ml chloramphenicol, and 0.04 mg/ml gentamicin (SGA, Sigma-Aldrich), potato glucose agar with 25 µg/ml chloramphenicol (PDA, Sigma-Aldrich), and malt extract agar with 0.1 mg/ml chloramphenicol (MEA, Sigma-Aldrich) (Supplementary Table 2). Plates were inoculated at 25 °C for 7 days.

After incubation, colonies were harvested with 1 ml PBS. Colonies in PBS were homogenized then divided in two: 500 µl was set aside for DNA extraction; BHI with 50% glycerol was added to the remaining for subsequent cultures. For bacteria, two sets of extraction were done per sample under 5% CO_2_ or anaerobic conditions. For fungi, samples were clustered in 12 groups according to age-sex-spirometry (Supplementary Table 1). Bacterial DNA extractions were conducted with Wizard® Genomic DNA Purification Kit (Promega) and fungal DNA extractions with Quick-DNA Fungal/Bacterial MiniPrep Kit (Zymo Research). Next generation sequencing libraries were carried out using the Nextera DNA Flex Library Prep kit (Illumina) following the manufacturer’s instructions. Paired-end metagenomic sequencing was performed on the Illumina Novaseq 6000 platform, generating 2×250 bp paired-end reads (Next-generation sequencing platform, CR-CHU de Québec-Université Laval). The mean number of reads per sample for bacteria were 57,590,398 reads and per group for fungus 268,625,462 reads.

### Bioinformatic analysis of the sequencing from culture enrichment

Reads were trimmed using Trimmomatic ^35^. For bacterial taxonomic profiling, sequence reads were mapped against the MetaPhlAn4 clade-specific marker gene database mpa_vJun23_CHOCOPhlAnSGB_202403 ^36^. For the culture-enriched draft genomes (hereafter CEGs), the reconstruction workflow was analogous to that used for metagenome-assembled genomes (MAGs) ^1^. The assembling was executed with MEGAHIT v1.2.9 ^37^, the binning with bbmap v38.86 ^38^, CEGs with more than 95% completeness and less than 5% contamination were selected using CheckM v1.1.3 ^39^, the de-replication was performed with dRep v3.4.3 ^40^. The bacterial classification was achieved against database release 220 from GTDB-Tk v2.4.0 ^41^. The fungal classification was conducted using Kraken2 v2.1.3 and its PlusPF database (Standard plus Refseq protozoa & fungi). Chi-square tests were performed using RStudio (version 4.3.0) with the chisq.test() function from the base R package.

### Gene specific primer design

To isolate colonies, we designed gene specific primers. Bins with the concerned genus and closely and distantly related species genomes were selected (accession numbers given in Supplementary Table 6). Genome annotations were performed using PROKKA v1.14.5 ^42^. With the R library orthologr v0.4.1, all the selected genomes were blasted against a reference genome, GCF_001064145.1. This analysis showed two distinct groups within our CEGs (Supplementary Figure 4). To go further in the primer design, we selected the group with the most prevalent species, recurring in 27 samples out of the 180. The best candidate gene coding for a hypothetical protein, JVOJ01000036.1_32, was screened within this group using a custom Python script.

### Whole genome sequencing and analysis of unnamed species

Samples with the concerned genus were plated onto CHOC and blood agar media then inoculated at 35 °C, 24h in 5% CO_2_ conditions and 48h in anaerobic conditions. Isolated colonies were tested by PCR using the following JVOJ01000036.1_32 gene specific primers: forward 5’-ATGAGCGYGCCRCTGCTRACGGTGAT-3’; reverse: 5’-AGTTGAAGCCGAGGACMGCCGCC-3’. DNA extractions of positive colonies were performed with Wizard® Genomic DNA Purification Kit (Promega). Next generation sequencing libraries were carried out using the Nextera DNA Flex Library Prep kit (Illumina) following the manufacturer’s instructions and sequenced using the Illumina NovaSeq 6000 technology (Next-generation sequencing platform, CR-CHU de Québec-Université Laval). The mean number of reads per genome was 4,287,873 reads for three isolates named 729a-P8-7, 729a-cl1, and 729a-cl2 and 40,787,547 reads for the other isolates. Reads were trimmed using Trimmomatic ^35^. The scaffolds construction was performed using spades v3.15.4 ^43^ with --isolate option, the read alignment using bbmap v39.06 ^38^. Contigs with less than 500 bp were excluded then the polishing step were performed using pilon v1.24 ^44^, the genome annotation with PROKKA v1.14.5 ^42^, the gene clustering using Panaroo v1.0.1 ^45^. The quality was checked with CheckM v1.1.3 ^39^, and QUAST v5.2.0 ^46^. In all cases, the reconstructed genomes exhibited comparable genome sizes and GC content. The multiple sequence alignment was performed with mafft v7.526 ^47^, the phylogenetic trees were constructed using FastTree v2.1 ^48^ and visualized using Dendroscope 3 v3.8.10 ^49^.

### Phenotypic descriptions of unnamed species

Gram colorations were performed. The bacteria were observed under a transmission electron microscope (Imaging - Microscopy Platform, IBIS, Université Laval, and Bioimaging platform, CR-CHU de Québec-Université Laval). Bacteria were fixed with 2.5% glutaraldehyde in 0.1 M cacodylate buffer, then colored 8 min with uranyl acetate and 3 min with lead citrate.

## Supporting information

Supplementary Table III

## Declarations

### Ethics approval and Consent to participate

This study was approved by the Research Ethical Committee of Laval University Hospital center in Quebec City with project number MP-20-2019-4110 pertaining to the 2017 Qanuilirpitaa? health survey.

### Consent for publication

The final version of the manuscript has been approved by all authors.

### Availability of data and materials

The next generation sequencing dataset that supports the findings of this study was used under copyright agreement and are available upon request. It is stored on the VALERIA data management platform of Université Laval ([VALERIA](https://valeria.science/accueil)) under controlled access. In accordance with the First Nations principles of ownership, control, access, and possession (OCAP®), the Nunavik Regional Board of Health and Social Services is the owner of the data and of the biological samples collected during the Qanuilirpitaa? 2017 health survey on behalf of the Inuit population of Nunavik. Any request for data access should be addressed to the Qanuilirpitaa? 2017 Data Management Committee that oversees the management of the survey data as well as its biological samples and digital data. This management framework is integral to the research partnership process and is intended to allow researchers to access and use the data in a manner that is respectful of all parties. Additional information on data ownership, management, and access, as well as the Data/Biological Samples Request and Analysis Proposal Application Form are available in the Methodological report of the Qanuilirpitaa? 2017 (see Appendix ‘Policy on the management of databases and biological samples’, Section 7, pages 14–16 (415–417) and Appendix B, pages 23–27 (424–428); https://nrbhss.ca/sites/default/files/health_surveys/A11991_RESI_Rapport_methodologique_EP4.pdf).

### Authors’ contributions

Performed experiment: MF, HG; Analyzed data: MF, PL, NPP, JB; Drafted the manuscript: MF, PL; Revised the manuscript: MF, PL, BP, FM, MO; Designed the study and provided funding: BP, FM, MO

## Acknowledgments

We are grateful to the 2017 Nunavik Inuit Health Survey - Qanuilirpitaa? (Q2017) participants, as well as to all our Nunavik partners (including the Q2017 Data management committee, Q2017 Steering committee, and the Nunavik Regional Board of Health and Social Services), the Institut National de Santé Publique du Québec (INSPQ), as well as all Inuit and non-Inuit investigators who have collaborated in the various steps of the project and provided their intellectual input. We thank Cynthia Brouillard, IUCPQ for help with sample selection; Natalia Poliakova, CRCHUQ for help with coordination and monitoring; and Pierre Ayotte, INSPQ for his advice with the overall project and Sandra Isabel for bacterial isolation, Adnane Sellam for encouraging us to work with fungus. This research was enabled in part by computing infrastructure provided by Alliance Canada (www.alliancecan.ca) and Compute Canada (www.computecanada.ca). This research was supported by the Sentinel North program of Université Laval funded Canada First Research Excellence Fund. This work was also supported by a Canadian Institutes of Health Research Foundation Grant to MO. M.O. holds a Canada Research Chair in Antimicrobial Resistance. The funders played no role in study design, data collection, analysis and interpretation of data, or the writing of this manuscript.

## References

1 Flahaut, M. et al. Distinctive features of the oropharyngeal microbiome in Inuit of Nunavik and correlations of mild to moderate bronchial obstruction with dysbiosis. Scientific Reports 3, doi:10.1038/s41598-023-43821-4 (2023).

2 Robert, P. et al. (Nunavik Regional Board of Health and Social Services & Institut National de santé publique Québec, Kuujjuaq, 2020).

3 Whelan, F. J. et al. The Loss of Topography in the Microbial Communities of the Upper Respiratory Tract in the Elderly. Ann Am Thorac Soc 11, 13–521,, doi:10.1513/AnnalsATS.201310-351OC (2014).

4 Martellacci, L. et al. Characterizing Peri-Implant and Sub-Gingival Microbiota through Culturomics. First Isolation of Some Species in the Oral Cavity. A Pilot Study. Pathogens 9, doi:10.3390/pathogens9050365 (2020).

5 Lagier, J.-C. et al. Microbial culturomics: paradigm shift in the human gut microbiome study. Clinical Microbiology and Infection 18, 1185–1193, doi:10.1111/1469-0691.12023 (2012).

6 Fleming, E. et al. Cultivation of common bacterial species and strains from human skin, oral, and gut microbiota. BMC Microbiology 21, doi:10.1186/s12866-021-02314-y (2021).

7 Martellacci, L. et al. A Litterature Review of Metagenomics and Culturomics of the Peri-implant Microbiome: Current Evidence and Future Perspectives. Materials (Basel) 12, doi:10.3390/ma12183010 (2019).

8 Khelaifia, S. et al. Culturing the Human Oral Microbiota, Updating Methodologies and Cultivation Techniques. Microorganisms 11, doi:10.3390/microorganisms11040836 (2023).

9 Bilen, M. et al. The contribution of culturomics to the repertoire of isolated human bacterial and archaeal species. Microbiome 6, doi:10.1186/s40168-018-0485-5 (2018).

10 Nouioui, I. et al. Genome-Based Taxonomic Classification of the Phylum Actinobacteria. Frontiers in Microbiology 22, doi:10.3389/fmicb.2018.02007 (2018).

11 Zhao, Y. et al. Exploration of lung mycobiome in the patients with non-small-cell lung cancer. BMC Microbiology 23, doi:10.1186/s12866-023-02790-4 (2023).

12 Duda, J. J. & Slack, J. M. Ultrastructural Studies on the Genus Actinomyces. Journal of General Microbiology 71, 63–68, doi:10.1016/0003-9969(74)90228-3 (1972).

13 Sun, Y. et al. Characterization of Lung and Oral Microbiomes in Lung Cancer Patients Using Culturomics and 16S rRNA Gene Sequencing. Microbiology Spectrum 11, 1–12, doi:10.1128/spectrum.00314-23 (2023).

14 Tiew, P. Y. et al. A high-risk airway mycobiome is associated with frequent exacerbation and mortality in COPD. European Respiratory Journal 57, doi:10.1183/13993003.02050-2020 (2021).

15 Binte Mohamed Ali, N. A., et al. The Healthy Airway Mycobiome in Individuals of Asian Descent. CHEST 159, 544–548, doi:10.1016/j.chest.2020.09.072 (2021).

16 Rouhi, F. et al. Yeast species in the respiratory samples of COVID-19 patients; molecular tracking of Candida auris. Frontiers in Cellular and Infection Microbiology 14, doi:10.3389/fcimb.2024.1295841 (2024).

17 Chen, B.-Y. et al. Characteristics and Correlations of the Oral and Gut Fungal Microbiome with Hypertension. Microbiology Spectrum 11, doi:10.1128/spectrum.01956-22 (2023).

18 Xu, X. et al. Upper respiratory tract mycobiome alterations in different kinds of pulmonary disease. Frontiers in Microbiology 14, doi:10.3389/fmicb.2023.1117779 (2023).

19 Jean-Christophe Lagier, P. H., Saber Khelaifia, Pierre-Edouard Fournier, Bernard La Scola, Didier Raoult The rebirth of culture in microbiology through the example of Culturomics to study human gut microbiota. Clinical Microbiology Review 28, 237–264, doi:10.1128/CMR.00014-14 (2015).

20 Itagaki, T., Sakata, K.-I., Hasebe, A. & Kitagawa, Y. Exploratory Study of the Relationship between an Oral Fungal Swab Test and Patient Blood Test Data. Microorganisms 11, doi:10.3390/microorganisms11122887 (2023).

21 Khadija, B., Imran, M. & Faryal, R. Keystone salivary mycobiome in postpartum period in health and disease conditions. Journal of Medical Mycology 31, doi:10.1016/j.mycmed.2020.101101 (2021).

22 Angoulvant, A., Guitard, J. & Hennequin, C. Old and new pathogenic Nakaseomyces species: epidemiology, biology, identification, pathogenicity and antifungal resistance. FEMS Yeast Research 16, doi:10.1093/femsyr/fov114 (2016).

23 Wei, N. et al. Characterization of oral bacterial and fungal microbiome in recovered COVID-19 patients. BMC Microbiology 23, doi:10.1186/s12866-023-02872-3 (2023).

24 Gosse, L., Amrane, S., Mailhe, M., Dubourg, G. & Lagier, J.-L. Capnocytophaga sputigena: An unusual cause of community-acquired pneumonia. IDCases, doi:10.1016/j.idcr.2019.e00572 (2019).

25 Zhang, X., Li, C., Cao, W. & Zhang, Z. Alterations of Gastric Microbiota in Gastric Cancer and Precancerous Stages. Frontiers in Cellular and Infection Microbiology 11, doi:10.3389/fcimb.2021.559148 (2021).

26 Ju, Y. et al. Integrated large-scale metagenome assembly and multi-kingdom network analyses identify sex differences in the human nasal microbiome. Genome Biology 25, doi:10.1186/s13059-024-03389-2 (2024).

27 Zaura, E. et al. On the ecosystemic network of saliva in healthy young adults. International Society for Microbial Ecology 11, 1218–1231, doi:10.1038/ismej.2016.199 (2017).

28 Aida, M. et al. Heyndrickxia coagulans strain SANK70258 suppresses symptoms of upper respiratory tract infection via immune modulation: a randomized, double-blind, placebo-controlled, parallel-group, comparative study. Frontiers in Immunology 15, doi:10.3389/fimmu.2024.1389920 (2024).

29 Li, W. et al. A catalog of bacterial reference genomes from cultivated human oral bacteria. NPJ Biofilms Microbiomes 1, doi:10.1038/s41522-023-00414-3 (2023).

30 Cuthbertson, L. et al. Genomic attributes of airway commensal bacteria and mucosa. Communications Biology 7, doi:10.1038/s42003-024-05840-3 (2024).

31 Chorlton, S. D. Ten common issues with reference sequence databases and how to mitigate them. Frontiers in Bioinformatics 4, doi:10.3389/fbinf.2024.1278228 (2024).

32 He, J. et al. Association between oral microbiome and five types of respiratory infections: a two-sample Mendelian randomization study in east Asian population. Frontiers in Microbiology 10, doi:10.3389/fmicb.2024.1392473 (2024).

33 Jiang, W. et al. Investigating oral microbiome profiles in patients with cleft lip and palate compared with the healthy control. BMC Oral Health 24, doi:10.1186/s12903-024-04387-3 (2024).

34 Lyu, X., Xu, X., Shen, S. & Qin, F. Genetics causal analysis of oral microbiome on type 2 diabetes in East Asian populations: a bidirectional two-sample Mendelian randomized study. Frontiers in Endocrinology 15, doi:10.3389/fendo.2024.1452999 (2024).

35 Bolger, A. M., Lohse, M. & Usadel, B. Trimmomatic: a flexible trimmer for Illumina sequence data. Bioinformatics 30, 2114–2120, doi:10.1093/bioinformatics/btu170 (2014).

36 Blanco-Míguez, A. et al. Extending and improving metagenomic taxonomic profiling with uncharacterized species with MetaPhlAn 4. Nature Biotechnology 41, 1633–1644, doi:10.1038/s41587-023-01688-w (2023).

37 Li, D., Liu, C.-M., Luo, R., Sadakane, K. & Lam, T.-W. MEGAHIT: an ultra-fast single-node solution for large and complex metagenomics assembly via succinct de Bruijn graph. Bioinformatics 31, 1674–1676, doi:10.1093/bioinformatics/btv033 (2015).

38 Bushnell, B., Rood, J. & Singer, E. BBMerge – Accurate paired shotgun read merging via overlap. PLoS ONE 12, 1–15, doi:10.1371/journal.pone.0185056 (2017).

39 Parks, D. H., Imelfort, M., Skennerton, C. T., Hugenholtz, P. & Tyson, G. T. Assessing the quality of microbial genomes recovered from isolates, single cells, and metagenomes. Genome Research 25, 1043–1055, doi:10.1101/gr.186072.114 (2014).

40 Olm, M. R., Brown, C. T., Brooks, B. & Banfield, J. F. dRep: a tool for fast and accurate genomic comparisons that enables improved genome recovery from metagenomes through de-replication. ISME Journal 11, 2864–2868, doi:10.1038/ismej.2017.126 (2017).

41 Chaumeil, P.-A., Mussig, A. J., Hugenholtz, P. & Parks, D. H. GTDB-Tk2: memory friendly classification with the genome taxonomy database. Bioinformatics 38, 5315–5316, doi:10.1093/bioinformatics/btac672 (2022).

42 Seemann, T. Prokka: rapid prokaryotic genome annotation. Bioinformatics 30, 2068–2069, doi:10.1093/bioinformatics/btu153 (2014).

43 Bankevich, A. et al. SPAdes: A New Genome Assembly Algorithm and Its Applications to Single-Cell Sequencing. Journal of Computational Biology 16, 455–477, doi:10.1089/cmb.2012.0021 (2012).

44 Walker, B. J. et al. Pilon: An Integrated Tool for Comprehensive Microbial Variant Detection and Genome Assembly Improvement. PLoS One 9, doi:10.1371/journal.pone.0112963 (2014).

45 Tonkin-Hill, G. et al. Producing polished prokaryotic pangenomes with the Panaroo pipeline. Genome Biology 21, doi:10.1186/s13059-020-02090-4 (2020).

46 Mikheenko, A., Prjibelski, A., Saveliev, V., Antipov, D. & Gurevich, A. Versatile genome assembly evaluation with QUAST-LG. Bioinformatics 34, il42-il50, doi:10.1093/bioinformatics/bty266 (2018).

47 Katoh, K. & Standley, D. M. MAFFT Multiple Sequence Alignment Software Version 7: Improvements in Performance and Usability. Molecular Biology and Evolution 30, 772–780, doi:10.1093/molbev/mst010 (2013).

48 Price, M. N., Dehal, P. S. & Arkin, A. P. FastTree: Computing Large Minimum Evolution Trees with Profiles instead of a Distance Matrix. Molecular Biology and Evolution 26, 1641–1650, doi:10.1093/molbev/msp077 (2009).

49 Huson, D. H. & Scornavacca, C. Dendroscope 3: an interactive tool for rooted phylogenetic trees and networks. Systematic Biology 61, 1061–1067, doi:10.1093/sysbio/sys062 (2012).

