## Supplementary figures for "Culturomics unveils species and expands bacterial and fungal diversity in Inuit oropharyngeal microbiota"

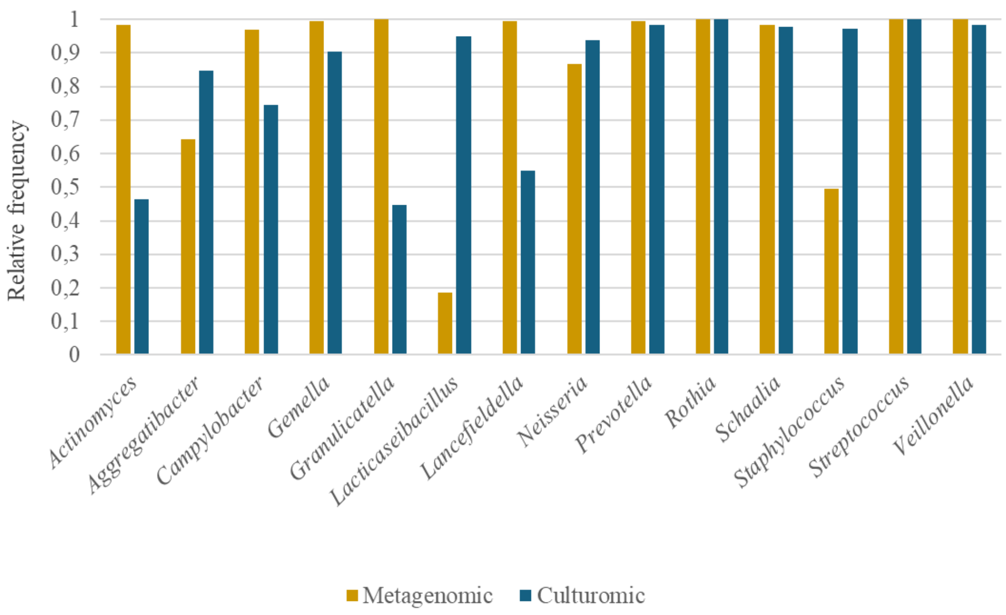


**Supplementary Figure 1**. The top ten most prevalent bacterial genera identified by MetaPhlAn4_202403 from the metagenomic and the culturomic datasets. The Y axis represents the proportion of samples that were positive for each genus by metagenomics (yellow) and culturomics (blue).

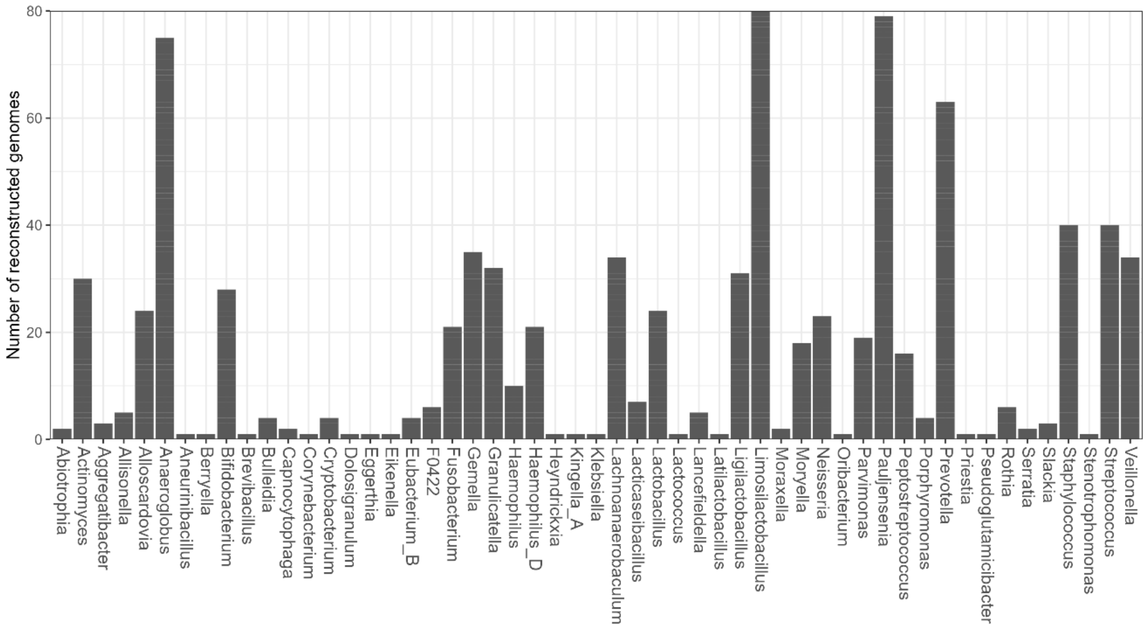


**Supplementary figure 2.** The number of reconstructed genomes per genus among our culturomic dataset.

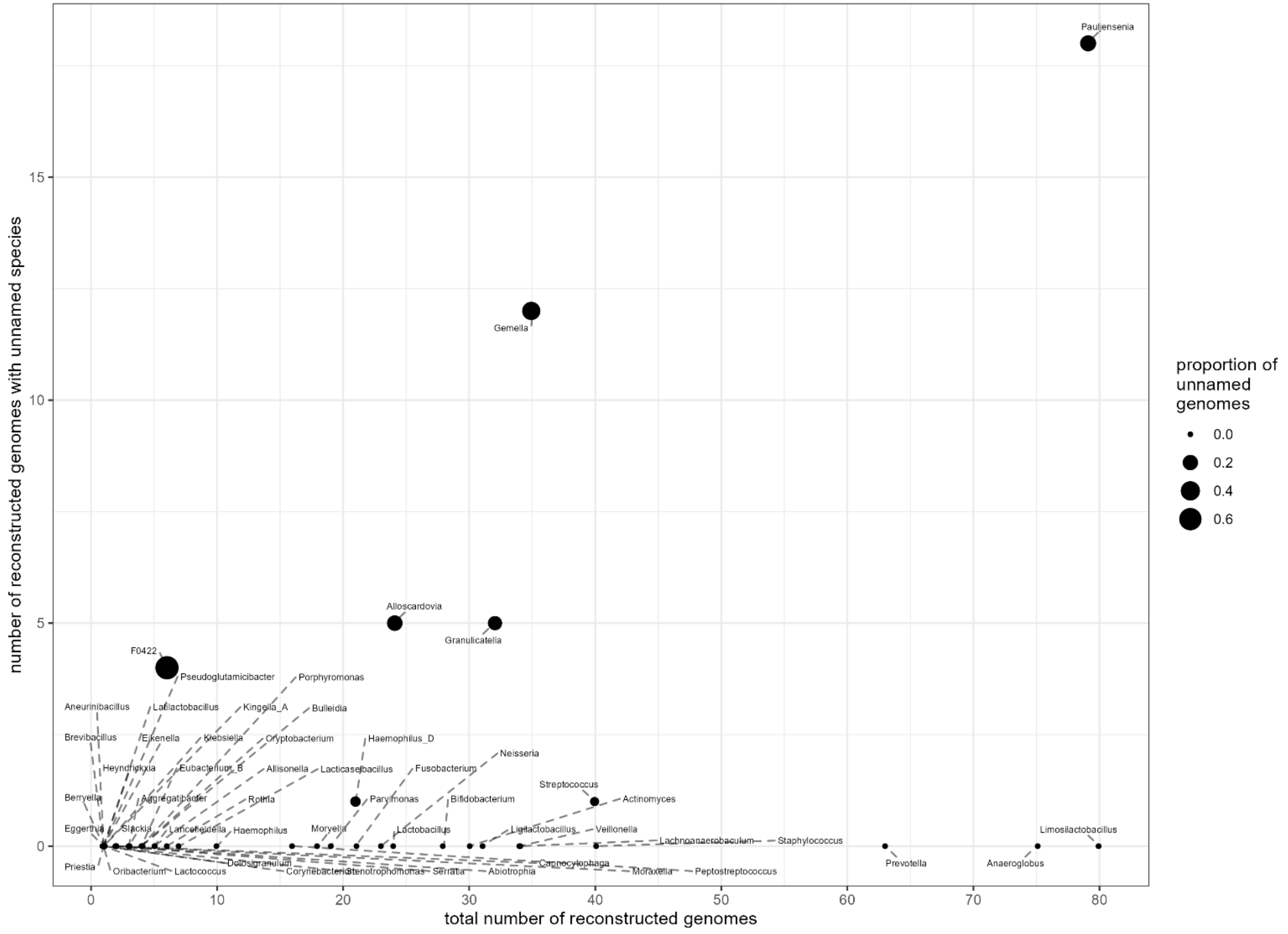


**Supplementary Figure 3.** The number of genomes with unnamed species compared to the total number of genomes per genus assembled from our culturomicsdataset.

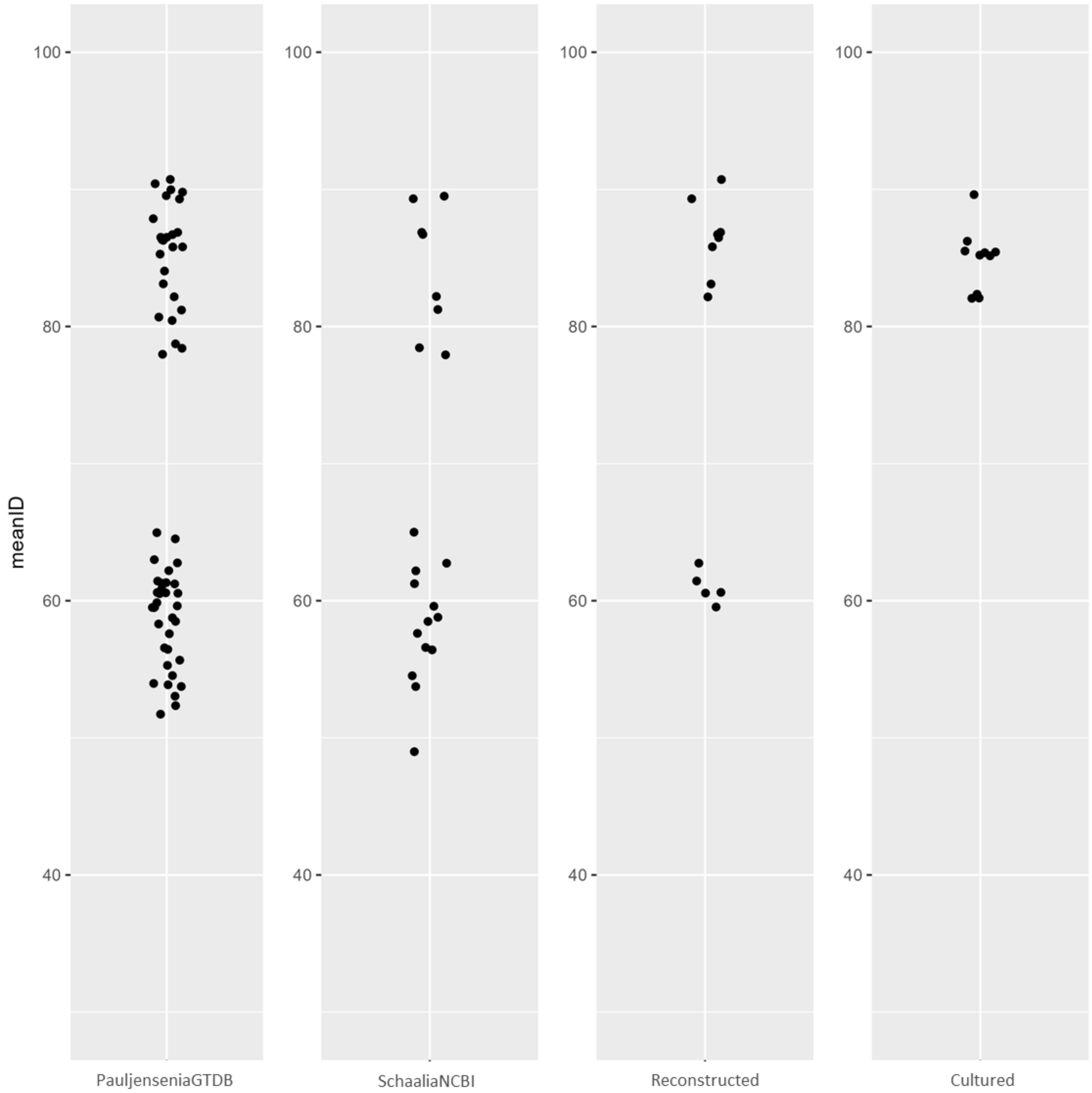


**Supplementary Figure 4.** Mean gene-level percent identities (computed from the entire set of ortholog genes per sample) for 104 genomes compared to *Pauljensenia* GCA_001064145.1. Each dot represents one of the 104 samples. PauljenseniaGTDB are the 60 *Pauljensenia* species genomes from GTDB v220; SchaaliaNCBI are the 21 NCBI type material referred as *Schaalia* species; Reconstructed are the culturomics genomes classified as *Pauljensenia* by GTDB-Tk v2.4.0; Cultured are our 10 *Pauljensenia* isolates recovered as single colonies on plates that were re-sequenced.

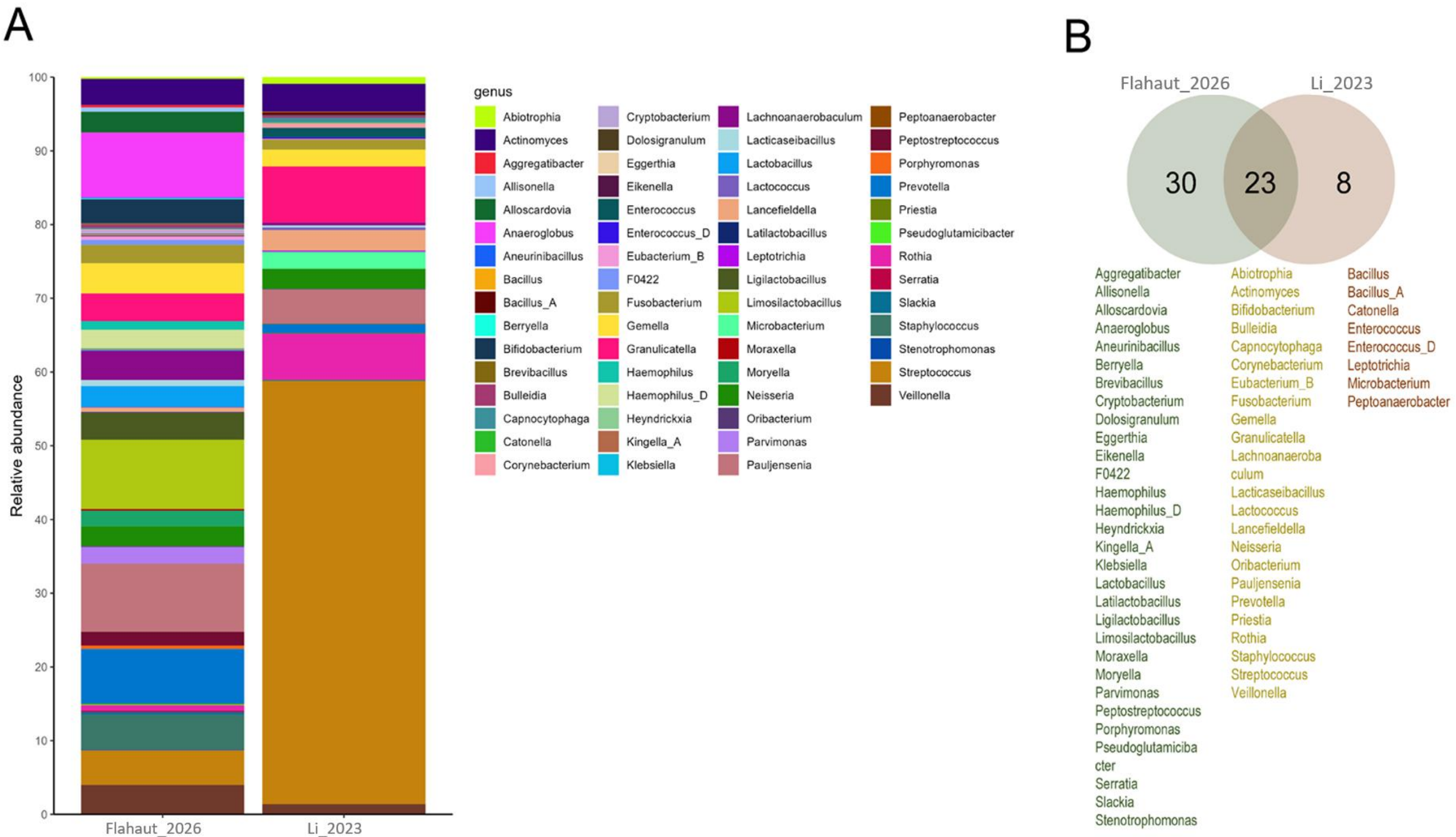
A
Relative abundance
0
10
20
30
40
50
60
70
80
90
100
Flahaut_2026
Li_2023
genus
Abiotrophia
Actinomyces
Aggregatibacter
Allisonella
Alloscardovia
Anaeroglobus
Aneurinibacillus
Bacillus
Bacillus_A
Berryella
Bifidobacterium
Brevibacillus
Bulleidia
Capnocytophaga
Catonella
Corynebacterium
Cryptobacterium
Dolosigranulum
Eggerthia
Eikenella
Enterococcus
Enterococcus_D
Eubacterium_B
F0422
Fusobacterium
Gemella
Granulicatella
Haemophilus
Haemophilus_D
Heyndrickxia
Kingella_A
Klebsiella
Lachnoanaerobaculum
Lacticaseibacillus
Lactobacillus
Lactococcus
Lancefieldella
Latilactobacillus
Leptotrichia
Ligilactobacillus
Limosilactobacillus
Microbacterium
Moraxella
Moryella
Neisseria
Oribacterium
Parvimonas
Pauljensenia
Peptoanaerobacter
Peptostreptococcus
Porphyromonas
Prevotella
Priestia
Pseudoglutamicibacter
Rothia
Serratia
Slackia
Staphylococcus
Stenotrophomonas
Streptococcus
Veillonella
B
Flahaut_2026
Li_2023
30
23
8
Aggregatibacter
Allisonella
Alloscardovia
Anaeroglobus
Aneurinibacillus
Berryella
Brevibacillus
Cryptobacterium
Dolosigranulum
Eggerthia
Eikenella
F0422
Haemophilus
Haemophilus_D
Heyndrickxia
Kingella_A
Klebsiella
Lactobacillus
Latilactobacillus
Ligilactobacillus
Limosilactobacillus
Moraxella
Moryella
Parvimonas
Peptostreptococcus
Porphyromonas
Pseudoglutamiciba
cter
Serratia
Slackia
Stenotrophomonas
Abiotrophia
Actinomyces
Bifidobacterium
Bulleidia
Capnocytophaga
Corynebacterium
Eubacterium_B
Fusobacterium
Gemella
Granulicatella
Lachnoanaeroba
culum
Lacticaseibacillus
Lactococcus
Lancefieldella
Neisseria
Oribacterium
Pauljensenia
Prevotella
Priestia
Rothia
Staphylococcus
Streptococcus
Veillonella
Bacillus
Bacillus_A
Catonella
Enterococcus
Enterococcus_D
Leptotrichia
Microbacterium
Peptoanaerobacter

**Supplementary Figure 5. (A)** The relative abundance of bacterial genera from the current study (Flahaut_2026) and from Li, *et al*., NPJ Biofilms and Microbiomes, 2023. **(B)** Venn diagram of the genera detected in the studies. The names of the genera from each category are indicated below the Venn diagram.

A.

```
>JVOJ01000036.1_32
ATGAGCGCGCCGCTGCTAACGGTGATCGTGCCCGCCTACAATTCCGAAGACTAC
CTTGATCGGGCGCTGACGACGCTCGTCGGATACGGCGACGAACTCGAAGCGATC
ATCGTCAACGACGGTTCGAAGGATCGCACCGCCCAGATTGCCGACGAGTGGGCC
GCCCGGTATCCCTCCGTGAGGGTTATCCACCAGGAAAACAAGGGCCATGGCGGC
GCCGTGAACGCCGGGCTCGCCGCCGCCACCGGTACGCACGTGCGCGTCGTTGAT
TCCGATGACTGGCTGGACCGACGCGCAACGAACGCCGTTCTCGACGTCCTGCGC
GAGGAACGCGAGGCCGGGCGCGACCTGGACCTGCTCGTCACCAACTACGTCTAC
GACAAGCAGGGAAAGTCGGTCAAGGCCGTCATCCGCTACCGCAACGTTCTGCCT
CGCGGACGTACCTTCGGCTGGGCCGACCTGCGCCGATGCCGCTACGACCAGTAC
CTGATGATGCACGCTCTGACGATGCGCACCGAGGTCGTGCGCGCGTCCGGCCTG
GTCATGCCCGAGCACACGTTCTACGTGGACTATCTCTACTCGTTCGTGCCGCTG
CCCTACATTTCGACGATCCGCTACCTGGACGTGGACCTGTACCACTACTTCATC
GGCCGCGACGACCAGTCCGTCAATGAGAAGGTCATGATCACGCGCTTGGATCAG
CTCGCCAGGGTGAACGAGGCGATGACCCGTGCGCTGCCCCCGCGCGCCGAGGTC
GAGGACAAGCTGTGGCGCTACATGGTCCACTACCTGCGCATCAACGCCGTCGCC
TGCTCCGTCATGGCTCAGCTCTCGGGCACCCCCGAGCACCTGGCCCTCAAGGAA
CAGATCTGGGAGACCATGGATCAGATCAACCCCGAGGCCACGGACCGCCTGCGC
CAGGACCTGCTCGCCGGCCTCGTGCGCCACGCGTCGCCGACGGTTGTTCGCGGC
GGCTACAAGGTCGCGGCGGCGGTCCTCGGCTTCAACTAA
```

B.

| Presence of JVOJ01000036.1_32 | Absence of JVOJ01000036.1_32 |
|---|---|
| GCF_001064145.1 | GCF_903645355.1 |
| GCF_900445025.1 | GCF_900499005.1 |
| GCF_900128465.1 | GCF_900155605.1 |
| GCF_900105015.1 | GCF_900155595.1 |
| GCF_024584435.1 | GCF_900155435.1 |
| GCF_019429565.1 | GCF_900106055.1 |
| GCF_009730335.1 | GCF_021083645.1 |
| GCF_005696695.1 | GCF_018986735.1 |
| GCF_002847525.1 | GCF_016405145.1 |
| GCF_000466265.1 | GCF_014208035.1 |
| GCF_000278725.1 | GCF_011038885.2 |
| GCA_916439125.1 | GCF_003858455.1 |
| GCA_916438945.1 | GCF_001746855.1 |
| GCA_916438365.1 | GCF_001070855.1 |
| GCA_902373545.1 | GCF_000820725.1 |
| GCA_902373435.1 | GCF_000758755.1 |
| GCA_900556405.1 | GCF_000429245.1 |
| GCA_019425105.1 | GCF_000429105.1 |
| GCA_018375675.1 | GCF_000420425.1 |
| GCA_001838165.1 | GCF_000308055.1 |
| GCA_001072465.1 | GCF_000296505.1 |
| GCA_000411415.1 | GCA_916719935.1 |
| | GCA_900554605.1 |
| | GCA_900541895.1 |
| | GCA_023426345.1 |
| | GCA_022649165.1 |
| | GCA_015655375.1 |

**Supplementary figure 6. (A)** Sequence of the gene JVOJ01000036.1_32 used for primer design targeting *Pauljensenia* sp. **(B)** Genomes from the GTDB database screened for the presence or absence of JVOJ01000036.1_32. Genomes containing JVOJ01000036.1_32 cluster in the upper group, whereas those lacking JVOJ01000036.1_32 cluster in the lower group (Supplementary Figure 4).
